# A compound PCP scheme underlies sequential rosettes-based cell intercalation

**DOI:** 10.1101/2022.11.09.515475

**Authors:** Yichi Xu, Yunsheng Cheng, Allison T. Chen, Zhirong Bao

**Affiliations:** Developmental Biology Program, Sloan Kettering Institute, New York, NY 10065, USA

## Abstract

Sequential rosettes are a type of collective cell behavior recently discovered in the *C. elegans* embryo to mediate directional cell migration through sequential formation and resolution of multicellular rosettes involving the migrating cell and its neighboring cells along the way. Here we show that a Planar Cell Polarity (PCP)-related polarity scheme regulates sequential rosettes, which is distinct from the known mode of how PCP regulates multicellular rosettes in the process of convergent extension. Specifically, non-muscle myosin (NMY) localization and edge contraction are perpendicular to that of Van Gogh as opposed to colocalizing with Van Gogh. Further analyses suggest a two-component polarity scheme: one being the canonical PCP pathway with MIG-1/Frizzled and VANG-1/Van Gogh localized to the vertical edges, the other being MIG-1/Frizzled and NMY-2 localized to the midline/contracting edges. Both MIG-1/Frizzled and VANG-1/Van Gogh are required for NMY-2 localization and contraction of the midline edges, but in redundancy with LAT-1/Latrophilin, an adhesion GPCR which has not been shown to regulate multicellular rosettes. Our results establish a distinct mode of PCP-mediated cell intercalation and shed light on the versatile nature of PCP pathway.

## INTRODUCTION

The Planar Cell Polarity (PCP) is a conserved mechanism to coordinate polarity of individual cells across a tissue (Butler and Wallingford, 2017). The coordination is controlled by the asymmetric distribution of core PCP components, Van Gogh and Frizzled. These two transmembrane proteins inhibit each other intracellularly to establish polarity within a cell while attracting each other intercellularly to synchronize the orientation of polarity between cells, thus forming a polarity that is capable to propagate to many cells (Amonlirdviman et al., 2005).

While such spatial propagation of polarity is most intuitive in static tissues, studies have shown that the PCP pathway can regulate directional cell movement to achieve global tissue shape change, such as convergent extension in vertebrate gastrulation, neurulation and organogenesis (Butler and Wallingford, 2017), as well as the assembly of the *C. elegans* central nerve cord neurons (Shah et al., 2017a). In these cases, the PCP pathway regulates cell intercalation through directional formation and resolution of multicellular rosettes (Lienkamp et al., 2012), where polarized localization of the PCP core components recruits non-muscle myosin to drive directional edge contraction. However, the PCP pathway is not the only mechanism to regulate directional formation and resolution of multicellular rosettes and tissue elongation. During Drosophila germband elongation, where directional formation and resolution of multicellular rosettes was first discovered (Blankenship et al., 2006), the process is driven by a planar polarized distribution of Toll-like receptors (Paré et al., 2014).

The core PCP pathway is known to interact with other pathways and genes in regulating tissue polarity. In the Drosophila wing disc and eye, protocadherins Fat and Dachsous show planar polarization, which can contribute to the planar polarization of the core pathway (Ambegaonkar et al., 2012; Brittle et al., 2012). During the elongation of *C. elegans* ventral nerve cord, SAX-3/Robo acts in parallel to VANG-1/Van Gogh (Shah et al., 2017a). A common feature is that these pathways regulate non-muscle myosin localization (Soto, 2017).

Recently, we identified a collective cell behavior in the *C. elegans* embryo, termed sequential rosettes, which mediates directional cell migration through sequential formation and resolution of multicellular rosettes involving the migrating cell and its neighboring cells along the way (Wang et al., 2022). In this study, we systematically examined cell movements in the early *C. elegans* embryo to show that sequential rosettes are a general mechanism mediating long-range cell migration in multiple cell lineages. Furthermore, we investigated the underlying molecular mechanism regulating sequential rosettes. We show that sequential rosettes present a novel mode of PCP-driven cell intercalation, where NMY-2 localization and edge contraction are orthogonal to the localization of the core PCP component Van Gogh. We term this mode PCP-ortho and propose a two-component polarity scheme: the canonical PCP pathway with MIG-1/Frizzled and VANG-1/Van Gogh localized to edges perpendicular to the contracting edges, and MIG-1/Frizzled, DSH-2/Disheveled and NMY-2 to the contracting edges. In addition, we identified LAT-1/Latrophilin, an adhesion-GPCR that is best known for regulating axon guidance and synapse formation (Moreno-Salinas et al., 2019), as a regulator of NMY localization and edge contraction in PCP-ortho and sequential rosettes.

## RESULTS

### Sequential rosettes mediate long-range cell migration in the *C. elegans* embryo

In the early *C. elegans* embryo, there are two major themes of cell movements that are intertwined, gastrulation and left-right patterning. Gastrulation, from the 26-cell stage to the 350-cell stage, involves in the internalization of many cells at multiple locations through the embryo (Harrell and Goldstein, 2011; Pohl et al., 2012). At the same time, left and right homologous lineages move to restore the superficial bilateral symmetry after the symmetry is broken at the 4-to-6 cell stage (Pohl and Bao, 2010; Pohl et al., 2012). Despite the technical capacity to image and track the position of every cell through this developmental time window, it is difficult to pinpoint individual cell movements due to whole embryo rotations (Moore et al., 2013).

To this end, we designed a computational method based on neighborhood change to identify migrating cells (Figure 1A, Methods). Briefly, the migration distance for each cell is defined as the average of the distances to initial neighbors when the cell is dividing and the distances to final neighbors when the cell is born. By focusing on relative movements, this approach avoids the complication of whole embryo rotations. In particular, this method is sensitive for leader cells in collective movement, which appear common during *C. elegans* embryogenesis.

**Figure 1.**
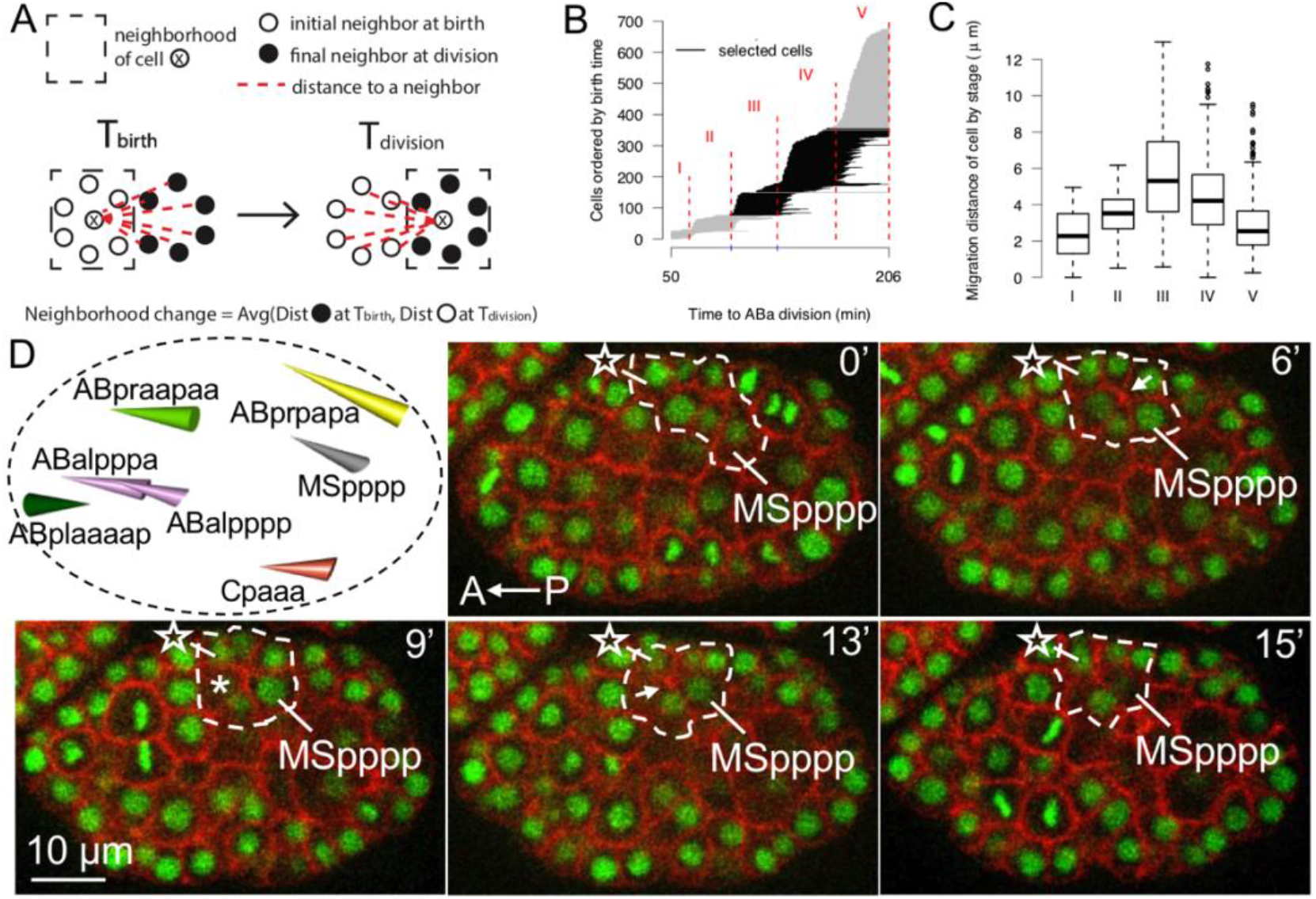
Sequential rosettes mediate long-range migration. (A) The computational screen of long-range migration by neighborhood change. (B) The birth and division timings of each cell from 1-cell to 350-cell stage. Each horizontal line denotes a cell starting at birth time and ending at division time sorted by birth time. Five stages were defined based on the timing of cell divisions (Methods). Black lines highlight Stages III and IV, which are of focus in this study. (C) The migration distance of cells in each stage from panel B. The center line denotes the median, while the box contains the 25th to 75th percentiles. The whiskers mark 1.5x interquartile range. (D)Top left panel, cone plot of the migration of 7 cells with sequential rosettes. The base of cone represents the position at birth of the cell and the vertex of cone represents the position at division of the cell. The color of cells denotes the founder lineage (see Figure S1 for convention). Fluorescent images show the sequential rosettes during the migration of MSpppp. The dash contour denotes cells involved in the sequential rosettes. The asterisk denotes the most anterior neighbor gained by MSpppp. The arrows at 6 and 13 mins denote the foci of rosettes. The asterisk at 9 min denotes new neighboring cell gained by MSpppp after the resolution of first rosette. Time 0 is 140 minutes after the diamond-shaped 4-cell stage. The embryo is ventral view and anterior to the left. Nuclei, HIS-72-GFP; plasma membrane, PH-domain of PLC1d1 fused to mCherry.

In wildtype embryos with complete lineage traced, we first divided early embryogenesis (1-to 350-cell stage, completion of gastrulation) into 5 stages based on synchronous cell divisions (Figure 1B, Methods). Stages III and IV, which involve 276 cells, show the most pronounced long migration distance (Figure 1C, Figure S1). In total, 37 cells were considered as migrating based on the distance (greater than mean + 1 standard deviation). Using images of embryos with fluorescently labeled nuclei and plasma membrane, we found that 18 of the 37 cells for which the movement is mediated by the T1 process of neighbor exchange and 12 by canonical multicellular rosettes (Figure S2). For the remaining 7 cells, the movement is mediated by sequential rosettes, each involving 2 or 3 consecutive rosettes. These cells are Cpaaa, ABalpppa, ABalpppp, ABplaaaap, ABpraapaa, ABprpapa, and MSpppp (Figure 1D). Cpaaa is the founding case of sequential rosettes that we previously identified (Wang et al., 2022). Furthermore, 4 of these cells (ABalpppa, ABalpppp, ABplaaaap, ABpraapaa) appear to be related to the restoration of bilateral symmetry involving the ABplaa/ABpraa lineages and their LR counterparts (Pohl and Bao, 2010). Combined with the migration of the mu_int_R cell, which is a case of long-range migration after gastrulation identified by Sulston (Sulston et al., 1983) and shown to be mediated by sequential rosettes in our previous study (Wang et al., 2022), these results suggest that sequential rosettes are a widely used mechanism of cell migration in the *C. elegans* embryo.

### Stereotypy of the Cpaaa sequential rosettes

We use the Cpaaa case to elucidate the regulatory mechanism of sequential rosettes. To better define the basis of cell behaviors for phenotype analyses, we first examined the dynamics of the Cpaaa sequential rosettes in wildtype embryos. The Cpaaa sequential rosettes involves 8 epidermal progenitor cells on the dorsal side of the *C. elegans* embryo (Wang et al., 2022). Around 170-200 mins after first cell cleavage, by sequential formation and resolution of rosettes, Cpaaa intercalates from posterior to anterior into two rows of three ABarpp cells (n1/n3/n5 and n2/n4/n6) and contacts ABarpaapp before cell division of Cpaaa (Figure 2A). After the resolution of each rosette, Cpaaa gains new neighbors, i.e., n3/4 at 6 min, n5/6 at 13 min, and ABarpaapp at 22 min (Figure 2A, asterisks and star cell). The intercalation of Cpaaa is essential for epidermis organization in *C. elegans*: the anterior and posterior chunk of the dorsal epidermis, represented by ABarpaapp and Cpaaa, respectively, get in contact and eventually fuse to form the large dorsal epidermal cell hyp7; the two rows of ABarpp cells will form the seam cells on the left and right side of the body.

**Figure 2.**
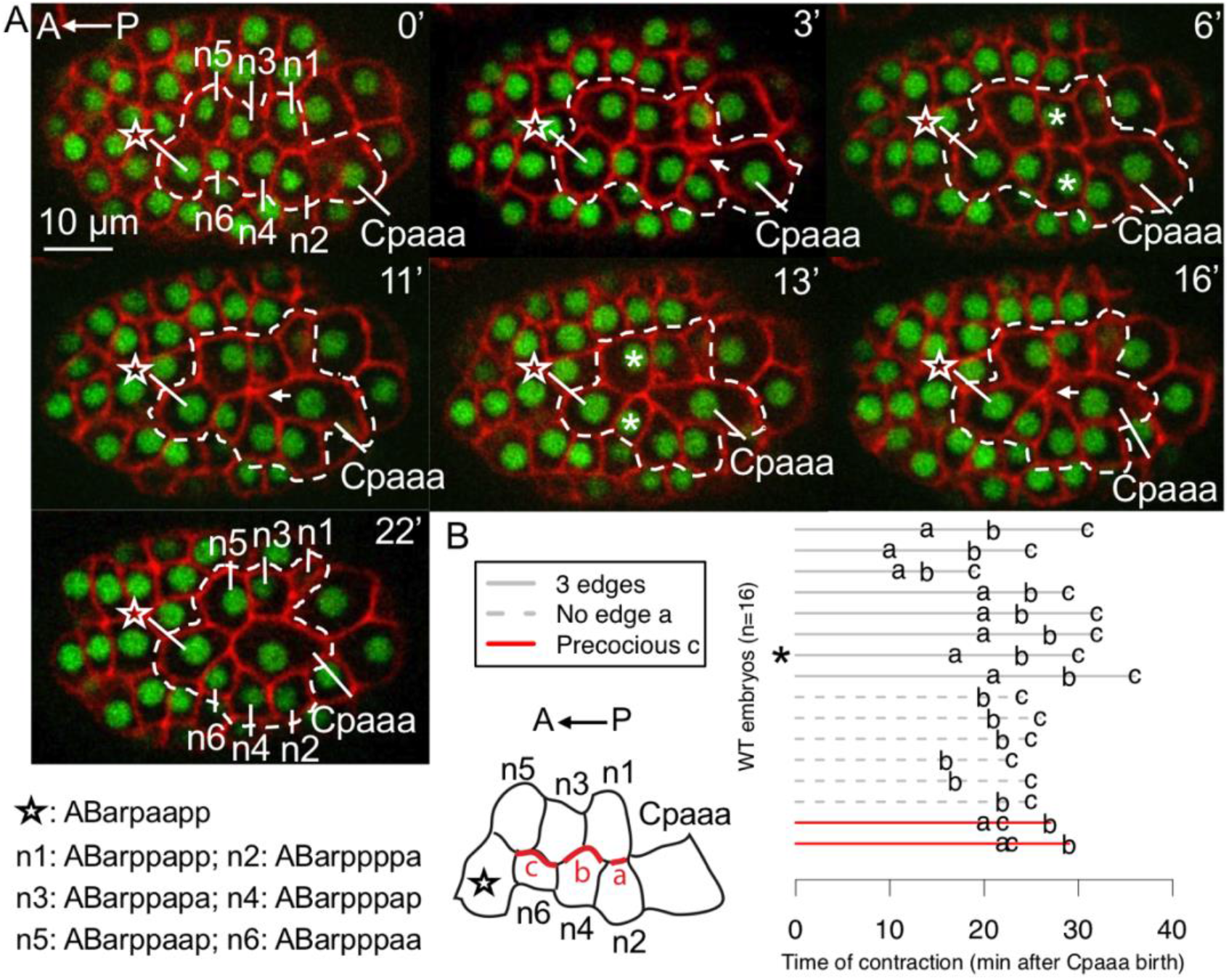
Sequential rosettes of Cpaaa. (A) Dynamics of rosette formation and resolution in sequential rosettes of Cpaaa. The contour denotes cells involved in the sequential rosettes. The star denotes the most anterior neighbor gained by Cpaaa. The arrows at 3, 11 and 16 mins denote the foci of rosettes. The asterisks at 6 and 13 min denotes new neighboring cells gained by Cpaaa after rosette resolution. The embryo is dorsal view and anterior to the left. For fluorescent labels, see Figure 1D. Time 0 is 1 minute before the formation of first rosette formation (about 150 minutes after the diamond-shaped 4-cell stage), which is applied to all images of Cpaaa rosette rosettes in this study. (B) The timings of contraction of edges in 16 wildtype embryos. Bottom left, three edges (a, b, and c) are defined from posterior to anterior in the typical order of contraction. Right panel, each line denotes an embryo and letters (a, b, c) denote the timing of contraction of the corresponding edge. The asterisk denotes the embryo showed in panel A. “Precocious c” means the contraction of edge c is earlier than the contraction of edge b. “No edge a” means no edge a in the initial conformation of cells.

The Cpaaa sequential rosettes occurs consistently in wildtype embryos, and Cpaaa always contacts ABarpaapp before its division (Figure 2B, N=16). However, there are variations in the initial conformations of these cells and the order of edge contractions. In the typical cases (8 out of 16), Cpaaa is born to contact n1 and n2; these cases involve three rosette formations and three edges contraction sequentially from posterior to anterior (namely, edge a, b, and c, Figure 2B). In 6 of 16 embryos, Cpaaa is born slighly more anteriorly and contacts n4; these cases involve two rosette formations and two edge contractions (Figure 2B, “No edge a”). In the least frequent cases (2 out of 16), although there are three edges in the initial conformation, the most anterior edge (edge c) contracts before the contraction of the edge in the middle (edge b) (Figure 2B, “Precocious c”). Overall, sequential rosettes have a highly consistent outcome despite some variations on the initial conformation and the order of contractions.

### Core components of the PCP pathway regulate sequential rosettes

To elucidate the molecular mechanism of sequential rosettes, we first examined the core components of the PCP pathway, Frizzled and Van Gogh. Both *mig-1/*Frizzled and *vang-1/*Van Gogh have strong expression in cells involved in the Cpaaa sequential rosettes based on scRNA-seq data (Figure S3) (Packer et al., 2019). We found that loss of function mutations in *mig-1* and *vang-1* display defects on two aspects of sequential rosettes, i.e., edge contraction and resolution. Specifically, a contraction defect is defined as the failure on the contraction of at least one edge, resulting in Cpaaa not contacting ABarpaapp. A resolution defect is defined as the prolonged existence of rosettes, which lasts more than 10 mins instead of immediate resolution as in the wild type. In *mig-1*(*e1787*) mutants (Pan et al., 2006), 17% embryos show contraction defects on edge b or edge c (Figure 3A, Table 1, N=30) and 16% embryos show resolution defects (Figure 3B, Table 1, N=30). In *vang-1*(*ok1142*) mutants, 38% embryos show resolution defects but none of them shows contraction defect (Figure 3B, Table 1, N=37). Together, our results show that core components of the PCP pathway regulate sequential rosettes of Cpaaa.

**Table 1.** Phenotypes of the Cpaaa sequential rosettes in *mig-1(e1787), vang-1(ok1142)* and *lat-1(RNAi)*. Numbers in parenthesis indicate total number of embryos for counting penetrance of each phenotype. For contraction defects (3rd column), only embryos without defective rosettes (5th column) were considered. For resolution defects (4th column), only embryos without contraction phenotype (3rd column) or defective rosettes (5th column) were considered. Defective rosettes in the 5th column means rosettes with additional endoderm cells but not Cpaaa (see section titled “Genetic interactions of *mig-1, vang-1*, and *lat-1*”).

| Mutant or RNAi | Number of embryos assayed | Number of embryos with contraction defect | Number of embryos with resolution defect | Number of embryos with defective rosettes |
| --- | --- | --- | --- | --- |
| *mig-1* | 30 | 5 (/30) | 4 (/25) | 0 (/30) |
| *vang-1* | 37 | 0 (/37) | 14 (/37) | 0 (/37) |
| *lat-1(RNAi)* | 34 | 22 (/34) | 0 (/12) | 0 (/34) |
| *mig-1, vang-1* | 32 | 5 (/26) | 10 (/21) | 6 (/32) |
| *mig-1, lat-1(RNAi)* | 32 | 18 (/32) | 2 (/14) | 0 (/32) |
| *vang-1, lat-1(RNAi)* | 30 | 16 (/30) | 5 (/14) | 0 (/30) |
| *mig-1, vang-1, lat-1(RNAi)* | 34 | 24 (/27) | 1 (/3) | 7 (/34) |

**Figure 3.**
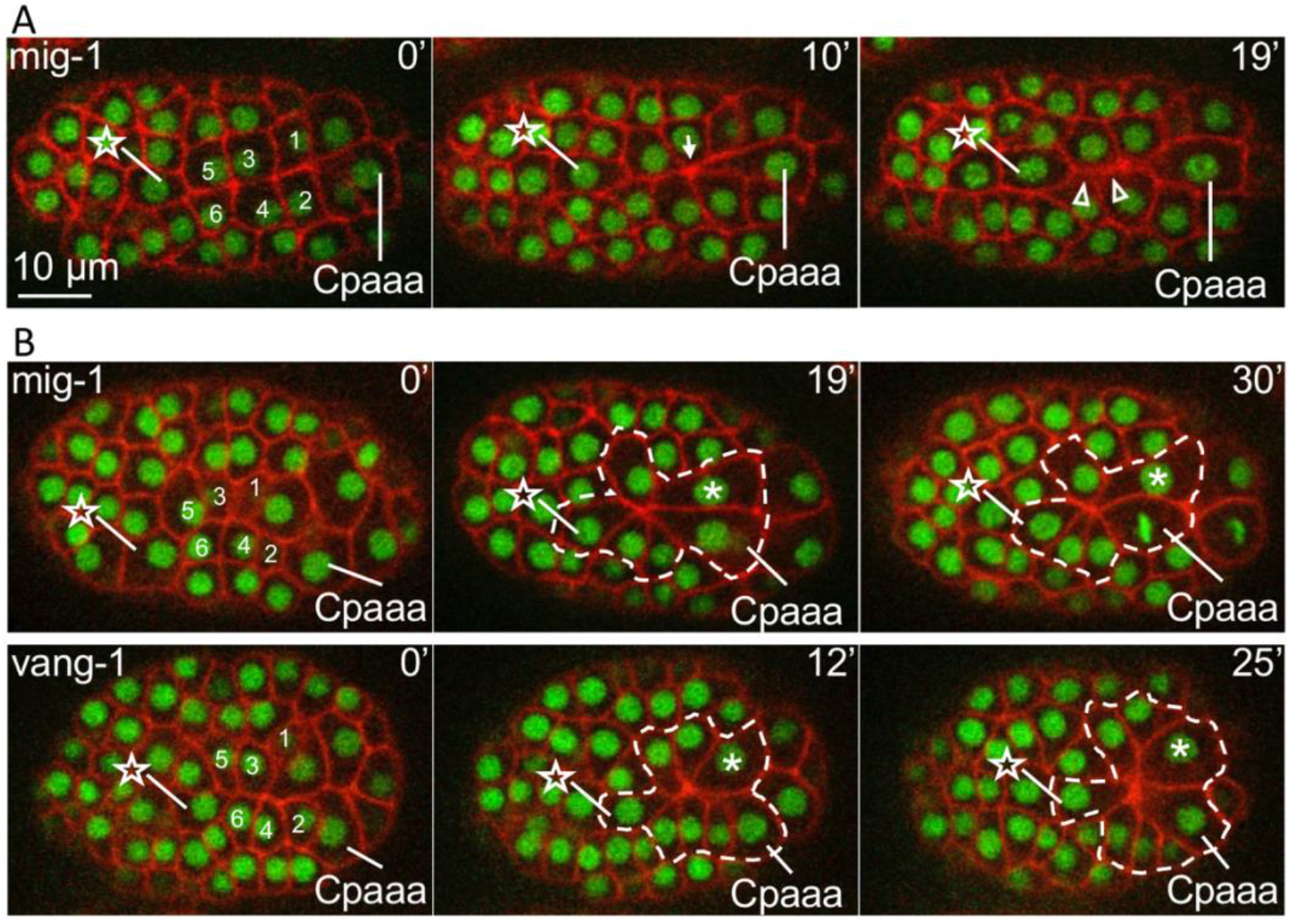
Core PCP genes regulate the sequential rosettes of Cpaaa. (A) Edge contraction defect in *mig-1(e1787)* mutant. The star and numbers denote 7 ABarp cells involved in the sequential rosettes (see Figure 2). The arrow at 10 min denotes the focus of first rosette. The arrowheads at 19 min denote edges (b and c) with contraction defect. (B) Rosette resolution defects in *mig-1(e1787)* (top) and *vang-1(ok1142)* (bottom) mutants. The contour denotes cells in the prolonged rosette. The asterisks denote Caaaa in the prolonged rosette, which participates in the first rosette in some of the wild type embryos. All embryos are dorsal view and anterior to the left. In all embryos, nuclei are marked by green channel and plasma membrane is marked by red channel as defined in Figure 1D.

We further examined other candidate genes. SAX-3/Robo is known to regulate multicellular rosettes in parallel to the PCP pathway in *C. elegans* during VNC convergent extension (Shah et al., 2017a). The *sax-3(ky123)* mutant, which affects rosettes in the VNC, did not show defects in the Cpaaa sequential rosettes (N=11). VAB-1 is an Eph receptor regulating the morphogenesis of epidermal cells at a later stage of *C. elegans* embryogenesis (Ghenea et al., 2005). *vab-1(dx31)* did not affect the Cpaaa sequential rosettes (N=11). Finally, Toll-like receptors regulate multicellular rosettes during Drosophila germband extension (Paré et al., 2014). *tol-1* is the sole homolog in *C. elegans*, but a loss of function allele, *nr2013*, did not show any defect in the Cpaaa sequential rosettes (N=13).

### MIG-1/Frizzled and VANG-1/Van Gogh localization indicate a PCP-related scheme

We then examined protein localization of MIG-1/Frizzled and VANG-1/Van Gogh. The contracting edges in the Cpaaa sequential rosettes coincide with the dorsal midline of the embryo and is parallel to the AP axis. For simplicity, we refer to these edges as the midline (red lines in the schematic in Figure 4A). The edges of the ABarpp cells that are approximately perpendicular to the midline/AP-axis are referred to as the vertical edges (green lines in the schematic in Figure 4A). The edges between the ABarpp cells and surrounding cells are referred to as lateral edges (grey lines in the schematic in Figure 4A). The midline and lateral edges collectively are referred to as horizontal edges.

**Figure 4.**
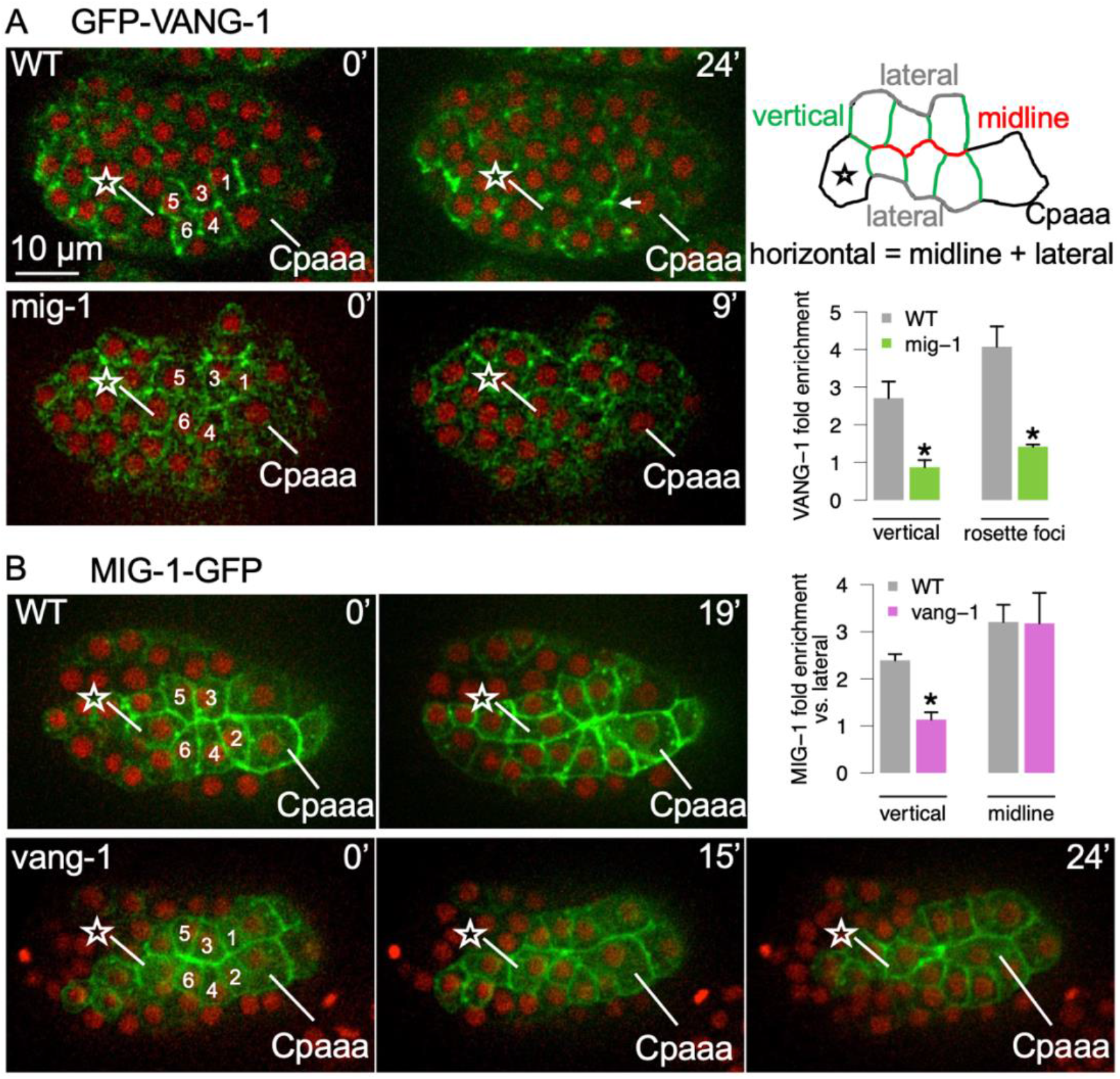
Protein localization of the core PCP components during the Cpaaa sequential rosettes. (A) GFP-VANG-1 localization in wildtype and *mig-1(e1787)* mutants. The arrow denotes rosette focus. The star and numbers denote ABarp cells (see Figure 2). The definitions of vertical, horizontal, lateral, and midline edges are shown in the diagram on initial conformation of cells before first contraction (top right). The bar plot shows VANG-1 fold enrichment between vertical and horizontal edges, and between rosette foci and edges in rosettes (bottom right). Error bar, standard error. *, Student’s t test p < 0.05. (B) MIG-1-GFP localization in wildtype and *vang-1(ok1142)* mutant. The bar plot shows MIG-1 fold enrichment between vertical and lateral edges, and between midline and lateral edges. See panel A for convention. All embryos are dorsal view and anterior to the left.

For VANG-1/Van Gogh, we used a CRISPR/Cas9-mediated GFP fusion at the N terminus of the native *vang-1* locus (*zy60*) (Shah et al., 2017a). Prior to the first edge contract, GFP-VANG-1 is enriched on the vertical edges, with an approximately 2.5-fold enrichment on the vertical edges versus horizontal edges (Figure 4A, 4 embryos, 4-6 cells per embryo, see Figure S4A for another example image). When rosettes form, GFP-VANG-1 is enriched at the foci, with an approximately 4-fold enrichment compared to the edges of the rosette (4 embryos, 4-6 cells per embryo). VANG-1 enrichment at rosette foci has been observed during the convergent extension of the *C. elegans* VNC (Shah et al., 2017a). Furthermore, VANG-1 localization to the vertical edges as well as rosette foci depends on *mig-1*. In *mig-1(e1787)* mutants, some ABarpp cells have VANG-1 localization on all edges, and foci enrichment is abolished (3 embryos, 4-6 cells per embryo).

For MIG-1/Frizzled, we used a transgenic line expressing a functional MIG-1-GFP fusion (Ex[Pmig-1::MIG-1::GFP]) (Mizumoto and Shen, 2013). The MIG-1-GFP fusion rescues the *mig-1(e1787)* phenotypes and displays wild type-like sequential rosettes (N=30). Similar to GFP-VANG-1, MIG-1-GFP is enriched on vertical edges (with an approximately 2.5-fold enrichment, 3 embryos, 4-6 cells per embryo) (Figure 4B). Interestingly, MIG-1-GFP is also enriched on contracting edges/midline, with an approximately 3-fold enrichment (3 embryos, 4-6 cells per embryo). Proper localization of MIG-1 depends on *vang-1*. In *vang-1(ok1142)* mutants, MIG-1-GFP enrichment on the vertical edges is lost: signals are similar between vertical and lateral edges (3 embryos, 4-6 cells per embryo). In contrast, MIG-1-GFP enrichment on the contracting edges/midline is not affected by *vang-1(ok1142)*.

The enrichment of VANG-1 and MIG-1 on the vertical edges as well as the mutual dependence of this localization are consistent with the canonical PCP model. Due to limitations of the reporters, we could not determine on which side of the cell-cell contact are the localizations.

The enrichment of MIG-1-GFP on the contracting edges/midline is distinct from the canonical PCP model. To rule out potential artifacts of overexpression by the transgene, we created a CRISPR/Cas9-mediated GFP fusion at the C terminus of the native *mig-1* locus. Unfortunately, the signal is too weak to be detected by imaging. However, as shown in experiments below, the midline localization of MIG-1 is most likely real and functional.

### A non-canonical PCP underlies sequential rosettes

Next, we examined the localization and regulation of NMY-2/non-muscle Myosin II. Using a CRISPR/Cas9-mediated GFP fusion of the native *nmy-2* locus (*cp13*), we found that NMY-2-GFP shows an approximately 3-fold enrichment on the contracting edges/midline (Figure 5A). More specifically, despite the sequential contraction of edges, NMY-2-GFP is enriched on all midline edges before the first contraction. NMY-2 localization and the direction of edge contraction in sequential rosettes shows a critical distinction from that in the canonical PCP model such as the VNC convergent extension (Shah et al., 2017a). In both cases, VANG-1 is localized to edges that are vertical to the midline/AP axis. In sequential rosettes, NMY-2 localization and the direction of edge contraction are perpendicular to VANG-1 localization. In contrast, during VNC convergent extension NMY-2 co-localizes with VANG-1 to drive contraction of the vertical edges. Thus, sequential rosettes employ a related but distinct scheme of polarity than the canonical PCP.

**Figure 5.**
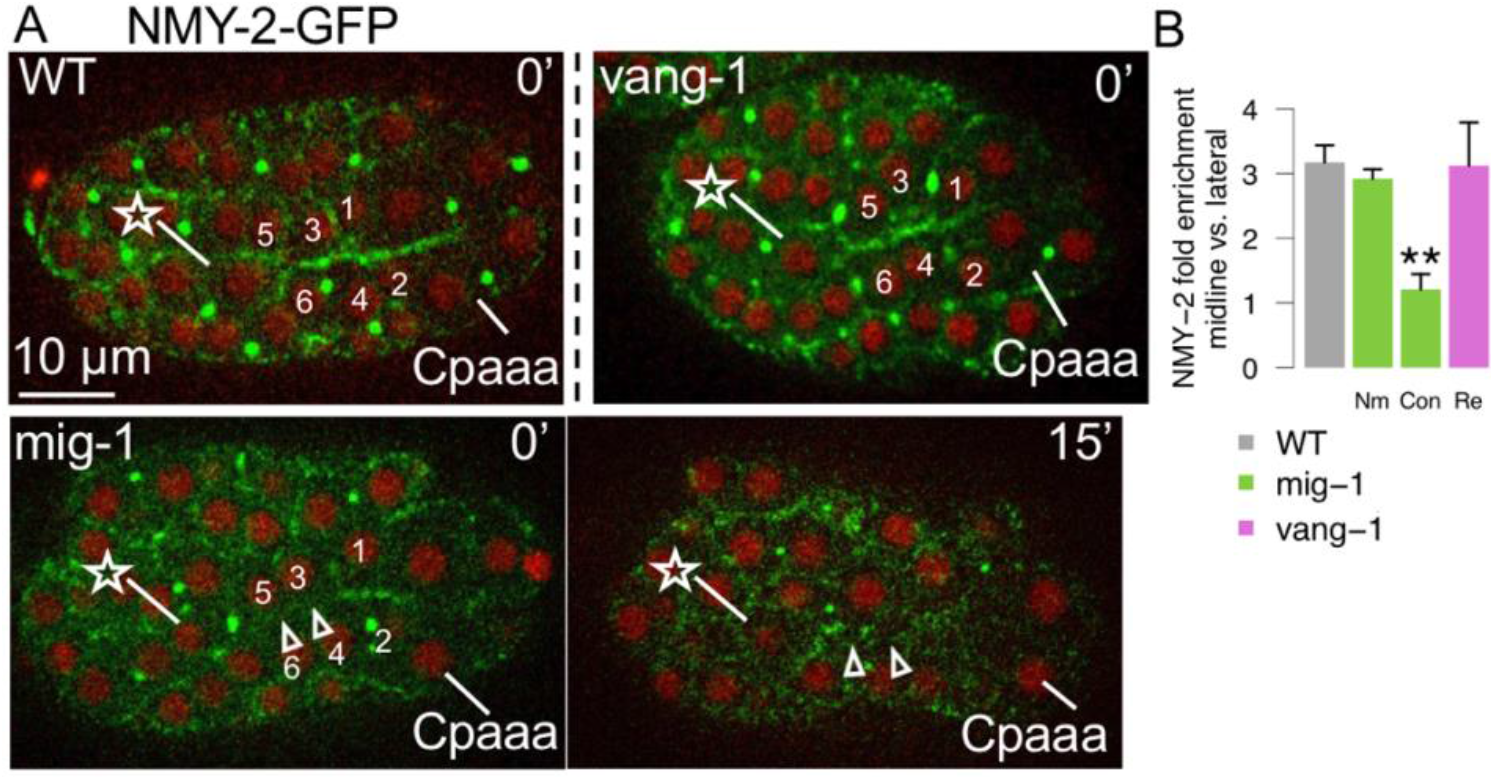
The regulation of midline NMY-2 localization in Cpaaa sequential rosettes. (A) NMY-2-GFP in sequential rosettes of Cpaaa in wildtype embryos, *mig-1(e1787)* and *vang-1(ok1142)* mutants. The arrows in *mig-1(e1787)* embryos denote edges with NMY-2 localization defect and contraction defect at 0 min and 15 min, respectively. The star and numbers denote ABarp cells (see Figure 2). All embryos are dorsal view and anterior to the left. (B) Bar plot shows NMY-2 fold enrichment between midline and lateral edges in wildtype and mutant embryos. Nm, edges with normal contraction on midline; Con, edges with contraction defect on midline; Re, only embryos with resolution phenotype were considered for *vang-1(ok1142)*. Error bar, standard error. **, Student’s t test p < 0.01.

We then asked how MIG-1/Frizzled and VANG-1/Van Gogh may regulate NMY-2 localization. In *mig-1(e1787)* mutants, enrichment of NMY-2 is lost on one or more midline edges in some embryos and the edges that lost NMY-2 enrichment displayed a contraction defect (Figure 5A, 7 embryos with contraction defects out of 32, comparable to the penetrance of edge contraction defects described above). In contrast, *vang-1(ok1142)* did not affect NMY-2 enrichment on the midline edges (Figure 5A, 0 out of 12 embryos), consistent with the lack of edge contraction phenotypes described above.

A minimal explanation of the phenotypic difference between *mig-1*/Frizzled and *vang-1*/Van Gogh loss of function is that midline localization of MIG-1/Frizzled is required for NMY-2 localization and edge contraction. Consistent with this explanation, we found that DSH-2/Dishevelled, a downstream effector that binds to Frizzled (Axelrod, 2001), is 2-fold enriched on the midline edges (Figure S4B). This observation is based on a CRISPR/Cas9-mediated mNeonGreen fusion of the native *dsh-2* locus at the N terminus (*cp51*), eliminating the concern of overexpression by traditional transgene reporters. Furthermore, *dsh-2* RNAi treated embryos displayed edge contraction defect with a penetrance (7 out of 38 embryos) similar to *mig-1(e1787)* (Figure S4C). These results suggest that midline enrichment of MIG-1/Frizzled is required for NMY-2 midline localization and edge contraction.

The results above indicate a two-component polarity scheme underlying sequential rosettes: a canonical PCP component with MIG-1/Frizzled and VANG-1/Van Gogh localization to the vertical edges with mutual dependence, and a midline component with NMY-2 and MIG-1/Frizzled localization on the midline edges. Given that NMY-2 localization and edge contraction are orthogonal to the edge with PCP protein localization, we term this polarity scheme PCP-ortho.

### LAT-1/Latrophilin is a regulator of PCP-ortho

Latrophilins are members of the adhesion G protein-coupled receptors (aGPCR) family, whose function is best known in the vertebrate nervous system for regulating synapse formation and function as well as actin dynamics in growth cones (Moreno-Salinas et al., 2019). The *C. elegans* homolog, LAT-1, is known to regulate spindle orientation in the early embryo (Langenhan et al., 2009), suggesting the potential to interact with Frizzled. Furthermore, *lat-1* is highly expressed in the ABarpp cells compared to others (Figure S3). Therefore, we asked whether *lat-1* could regulate sequential rosettes.

Loss of function of *lat-1* is maternal lethal, therefore we used RNAi to examine *lat-1* function in the Cpaaa sequential rosettes. We found that 65% embryos display the edge contraction defect and no embryo displays the rosette resolution defect (Figure 6A, Table 1, N=34). We then asked whether *lat-1* regulates edge contraction via NMY-2 localization. Indeed, the enrichment of NMY-2 on midline edges is abolished in *lat-1* RNAi treated embryos and the corresponding edges show contraction defects (Figure 6B, N=14). These results suggest that *lat-1* regulates NMY-2 localization and contraction of midline edges.

**Figure 6.**
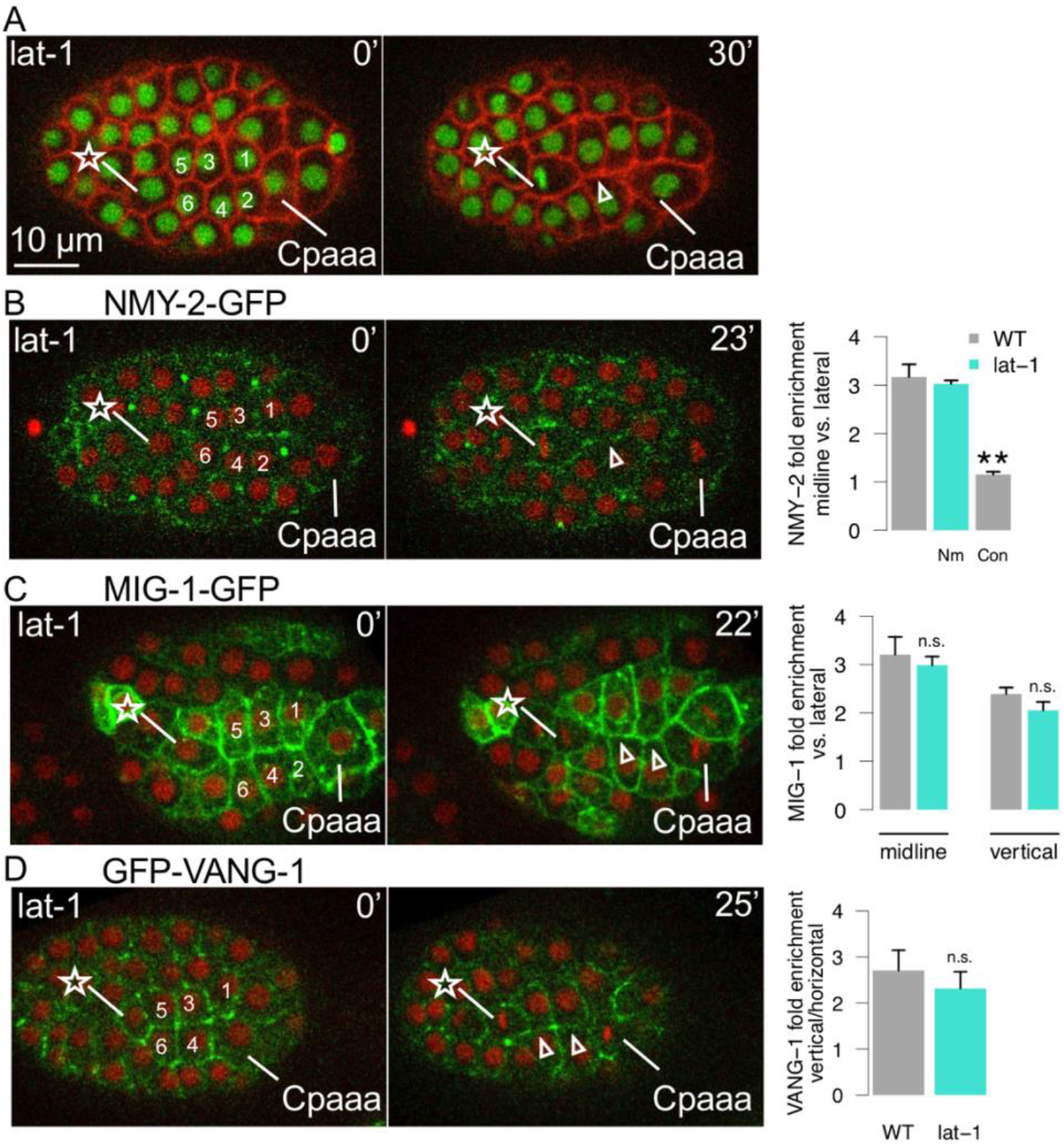
The regulation of *lat-1* in Cpaaa sequential rosettes. (A) The contraction defect (arrow) in embryos treated with *lat-1(RNAi)*. See Figure 2 for convention. (B) NMY-2-GFP on midline of sequential rosettes in wildtype embryos treated with *lat-1(RNAi)*. Nm, edges with normal contraction on midline; Con, edges with contraction defect on midline. (C) MIG-1-GFP on midline and vertical edges of sequential rosettes in wildtype embryos treated with *lat-1(RNAi)*. (D) GFP-VANG-1 on vertical edges of sequential rosettes in wildtype embryos treated with *lat-1(RNAi)*. The star and numbers denote ABarp cells (see Figure 2). Error bar, standard error. **, Student’s t test p < 0.01. All embryos are dorsal view and anterior to the left.

Next, we asked how *lat-1* may interact with *mig-1*/Frizzled and *vang-1*/Van Gogh. We first examined if *lat-1* is required for proper localization of MIG-1/Frizzled and VANG-1/Van Gogh. To account for the penetrance of *lat-1* RNAi knockdown, we focused on the embryos that showed edge contraction defects. We found the localization of MIG-1/Frizzled and VANG-1/Van Gogh is normal in these embryos (Figure 6C-D, N=10 and N=9, respectively): MIG-1/Frizzled is enriched on the vertical and midline edges and VANG-1/Van Gogh to the vertical edges, all with comparable fold of enrichment to the wild type. These results suggest that *lat-1* functions downstream of or in parallel with *mig-1*/Frizzled and *vang-1*/Van Gogh.

We then sought to examine the localization of LAT-1 by creating a CRISPR/Cas9-mediated GFP fusion at the C terminus of the native lat-1 locus. However, we were not able to generate a viable strain that can be stably maintained. Therefore, we focused the rest of the analyses on the phenotypes in double and triple loss of function of *lat-1(RNAi)* with *mig-1*/Frizzled and *vang-1*/Van Gogh.

### Genetic interactions of *mig-1, vang-1*, and *lat-1*

As described above, the three regulators cause two phenotypes in sequential rosettes, namely edge contraction and rosette resolution. We further found that *mig-1(e1787);vang-1(ok1142)* double mutants exhibit a third phenotype where the sequential rosettes involve additional cells from the endoderm (Figure 7, Table 1). In 6 out of 32 embryos, endoderm cells from E lineage, which normally lay underneath Cpaaa and the ABarp cells, were pulled by edge contraction to the dorsal and contact with ABarpaapp. It is known that *vang-1* is required for 8 endodermal cells (Exxx) at this stage to intercalate and form a plane (Asan et al., 2016). Further experiments are required to discern the tissue-specific gene functions and tissue interactions. Notably, in these embryos, Cpaaa disengages from the rosette after the initial participation. It is not clear if the disengagement is regulated, or a simple secondary consequence of excessive cell elongation and tension caused by the additional cells in the rosettes. Nonetheless, because Cpaaa no longer participates sequential rosettes with the ABarp cells in these embryos, we exclude them from further analysis (Table 1).

**Figure 7.**
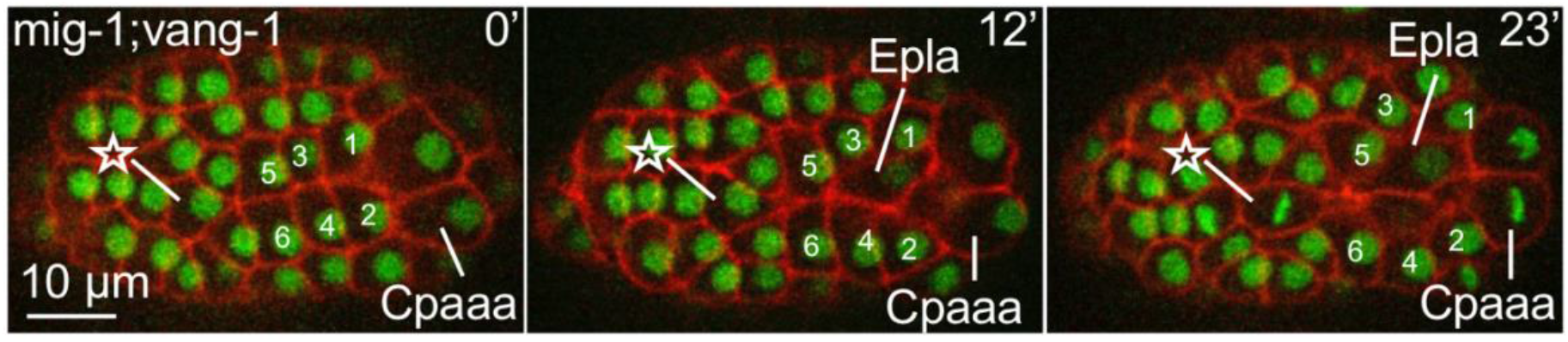
The rosettes involved in endoderm cell in *mig-1(e1787);vang-1(ok1142)* double mutant. Fluorescent images with dorsal view and anterior to the left. See Figure 2 for convention.

In terms of edge contraction, none of the three double loss of function conditions shows apparent enhancement compared to the single loss of function with stronger phenotype (Table 1 and Table S1, Fisher’s exact test p value > 0.05). However, the triple loss of function (*mig-1(e1787);vang-1(ok1142); lat-1(RNAi)*) shows significant increase of penetrance with nearly 90% of the embryos (24 out of 27, Table 1 and Table S1, Fisher’s exact p value 0.0087, 0.0041, and 2.9×10^−7^, respectively) showing edge contraction defects. The implication of the apparent three-way redundancy is discussed below.

Finally, in terms of rosette resolution, in *mig-1(e1787);vang-1(ok1142)* double mutants, 48% embryos show defects (Table 1), which is slightly higher than that in *vang-1(ok1142)* but not statistically significant (Table S1, Fisher’s exact test p value = 0.58). Furthermore, *lat-1(RNAi)* does not enhance the rosette resolution phenotype in *mig-1(e1787), vang-1(ok1142)* or *mig-1(e1787);vang-1(ok1142)* (Table 1 and Table S1, Fisher’s exact test p value = 1). These results suggest that *mig-1* and *vang-1* function in the same pathway in regulating rosette resolution while *lat-1* is not involved. In convergent extension, Van Gogh is required for proper rosette resolution (Williams et al., 2014). Our results extend the requirement to the core PCP pathway.

## DISCUSSION

### Diversity in PCP-mediated tissue rearrangement

This study revealed a mode of PCP-mediated tissue rearrangement that is distinct from the canonical convergent extension (Figure 8). Specifically, NMY localization and edge contraction occur at the edges orthogonal to VANG-1/Van Gogh localization. The resulting tissue shape change is widening and shortening along the AP axis, as opposed to the narrowing and lengthening in convergent extension. We term this mode PCP-ortho. Our results further suggested a polarity scheme with two components: the canonical Frizzled-Van Gogh localization to edges that are vertical to the AP axis, and Frizzled-NMY-2 localization to the midline/AP edges.

**Figure 8.**
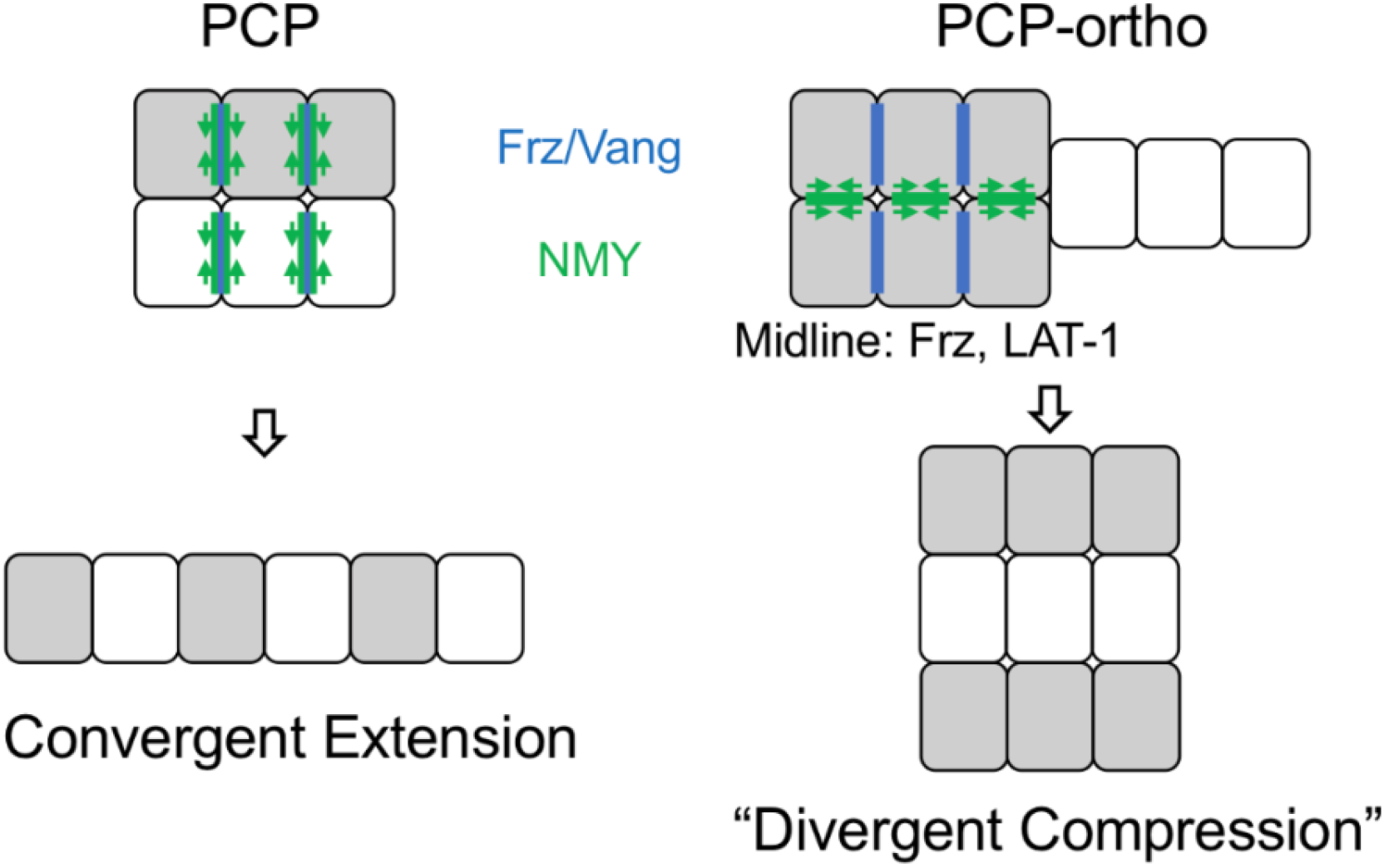
The comparison of canonical PCP model and PCP-ortho model.

PCP-ortho raises intriguing questions regarding cell polarity and regulation of NMY. In terms of NMY localization, both Frizzled and Van Gogh localization are known to recruit NMY in various contexts including convergent extension. What prevents NMY from following MIG-1/Frizzled or VANG-1/Van Gogh to the vertical edges in PCP-ortho? Furthermore, what recruits NMY-2 to the midline edges? Our results offer several clues. One is the population of MIG-1/Frizzled molecules on the midline edges. Given the differential localization of MIG-1/Frizzled and VANG-1/Van Gogh, it is possible that MIG-1/Frizzled and VANG-1/Van Gogh interaction on the vertical edges distinguishes the activity of MIG-1/Frizzled on the vertical vs midline edges. Another is the requirement of *vang-1* for midline localization of NMY-2, which was revealed in the triple loss of function *mig-1(e1787);vang-1(ok1142);lat-1(RNAi)*. This result indicates that signaling at the vertical edges acts not only *in cis* to repress NMY-2 localization to the vertical edges, but also *in trans* to promote NMY-2 localization to the midline edges. We hypothesize that the signaling at the vertical edges requires modification of the core Frizzled-Van Gogh pathway in canonical PCP. A slight complication is the apparent genetic redundancy of *mig-1*/Frizzled and *vang-1*/Van Gogh, which would typically argue against a single (linear) pathway. However, given the feedback loops in canonical PCP, the apparent redundancy may not be a concern. Clear elucidation of the polarity scheme would likely require optogenetic manipulation (Shah et al., 2017b) to separate the populations of molecules in different cells and on different edges.

The sequential rosettes also revealed nuanced temporal resolution of NMY-2 contraction activity. Our analysis showed that NMY-2 is localized to all midline edges prior to any edge contraction, while contraction occurs sequentially. These results indicate a latent signal to activate NMY-2. An apparent correlation with edge contraction is the contact with the Cpaaa cell, which could provide the latent signaling to activate NMY-2. As we previously showed, cell fate of Cpaaa is required for sequential rosettes (Wang et al., 2022).

### Flexibility of molecular interactions in the PCP pathway

Our study also reveals LAT-1/Latrophilin as an interactor with PCP pathway. The loss of function analyses suggest that *lat-1* is required for NMY-2 localization to the midline edges and that it acts in parallel to or downstream of *mig-1*/Frizzled and *vang-1*/Van Gogh. Further elucidation of *lat-1* function would benefit from knowing the localization of LAT-1.

The involvement of *lat-1* in a PCP scheme shows an intriguing similarity seen in the convergent extension of the *C. elegans* VNC neurons, where *sax-3*/Robo acts in parallel to *vang-1*. The PCP pathway relies on intricate feedback loops to synchronize polarity across a tissue. It is somewhat surprising that other polarity/NMY regulators can be so readily plugged into the scheme. In particular, both Latrophilin and Robo are known to function in adhesion, suggesting that their interactions with the PCP pathway could be intercellular to facilitate the synchronization of polarity across a tissue, similar to the protocadherins Fat and Dachsous in Drosophila (Ambegaonkar et al., 2012). Interestingly, many so-called axon guidance receptors are polarity/NMY regulators and are known to regulate cell intercalation in *C. elegans* (Asan et al., 2016; Bernadskaya et al., 2012; Chin-Sang et al., 1999). It is possible that these molecules can also be incorporated into a PCP context, either in *C. elegans* or in other organisms, which would greatly enrich our understanding of planar cell polarity.

In terms of the flexibility of the PCP pathway, previous studies in *C. elegans* indicated that even the so-called core interactions between Frizzled and Van Gogh may be partially replaced (Cravo and van den Heuvel, 2020). Regulation of spindle orientation in the early *C. elegans* embryo depends on polarized localization of MOM-5/Frizzled to the posterior of the dividing cell. Two features of MOM-5/Frizzled resembles planar cell polarity in that synchronized direction of polarity can arise among a group of cells independent of Wnt ligands (Park et al., 2004); and that the synchronization relies on cell-to-cell relay (Bischoff and Schnabel, 2006). *Vang-1* does not seem to be required in this process: while we did not characterize spindle orientation in our study, the fact that *vang-1* loss of function, either as single mutant or in combination with *mig-1*, is viable suggests that spindle orientation is largely not affected. Given that *vang-1* is the only apparent homolog of Van Gogh, it is likely that a different kind of membrane protein plays the role to interact with MOM-5/Frizzled to relay polarity, either on its own or redundant to *vang-1*. Lat-1/Latrophilin is known to regulate spindle orientation in the early embryo (Langenhan et al., 2009). It remains to be seen if and how the PCP-like scheme underlying spindle regulation is related to the PCP-ortho scheme underlying sequential rosettes.

## Supplementary Figures and Tables

**Figure S1.**
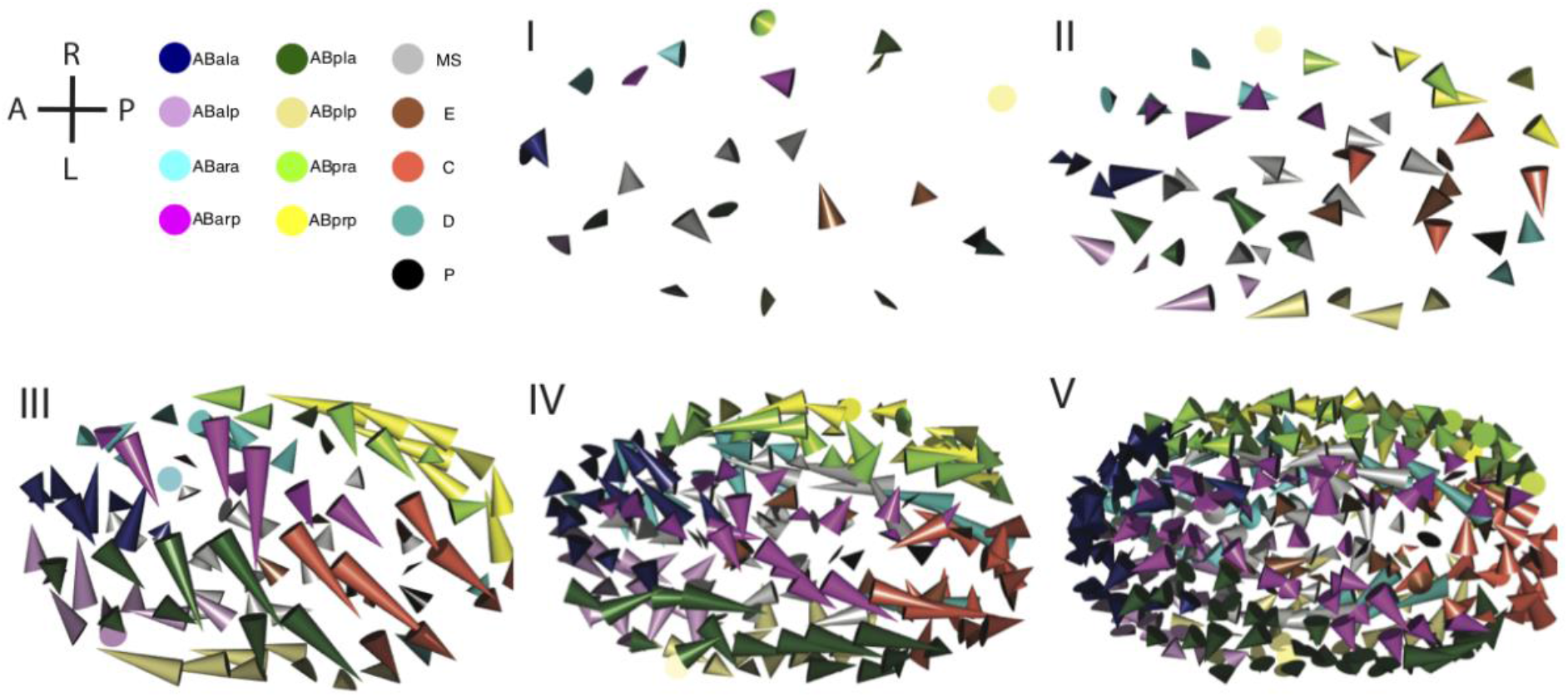
The migration of cell in 5 stages defined by cell divisions. Cone plots of the migration of cells in each stage from Figure 1B. The base of cone represents the position at birth of the cell and the vertex of cone represents the position at division of the cell. The color of cells denotes the founder lineage.

**Figure S2.**
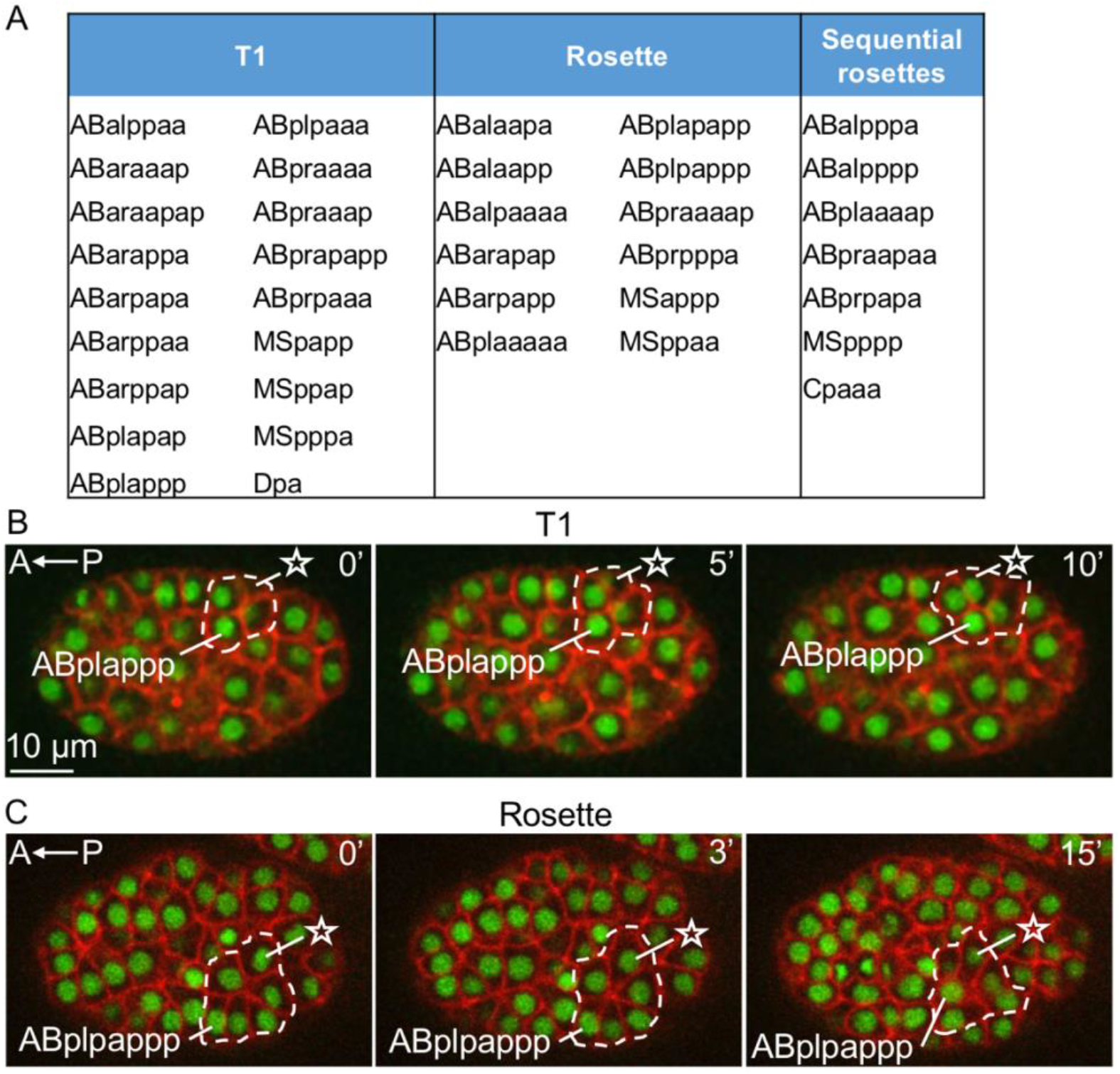
37 cells with large change of neighborhood. (A) Cells with T1, rosettes, and sequential rosettes, respectively. (B) Example of a cell with T1 process. Time 0 is 108 minutes after the diamond-shaped 4-cell stage. (C) Example of a cell with rosette. The star denotes the gained neighbor cell. Time 0 is 150 minutes after the diamond-shaped 4-cell stage. See Figure 1D for convention.

**Figure S3.**
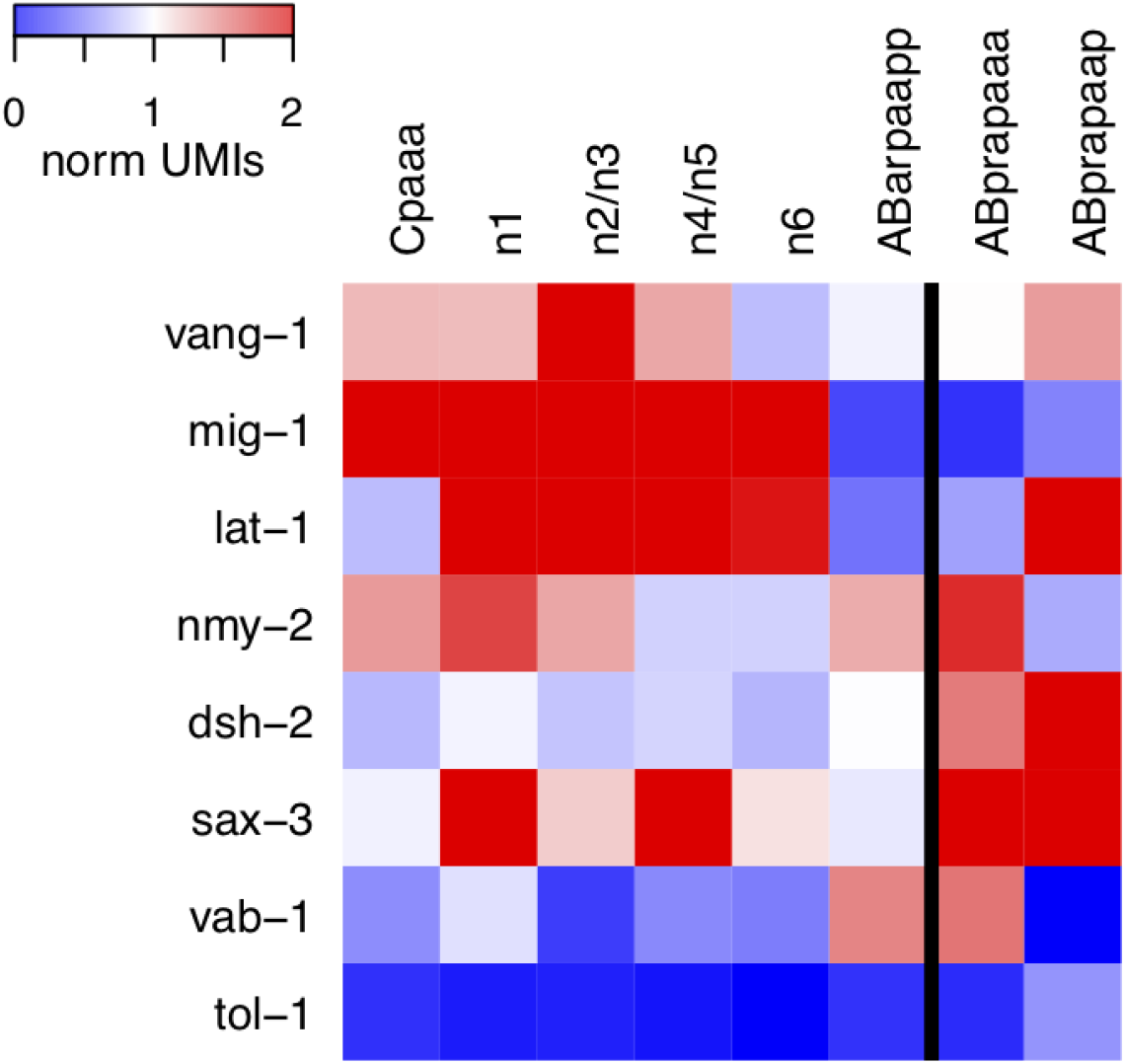
The expression of genes examined in this study in scRNA-seq data (Packer et al., 2019). The gene expression was normalized to UMIs per 10000 UMIs and calculated as mean of normalized UMIs in each cell. Cell annotation from the original publication was used. See Figure 2 for cell identity of n1-6. ABprapaaa and ABprapaap are showed as controls, which are two neighboring cells of ABarp cells but not involved in the sequential rosettes of Cpaaa.

**Figure S4.**
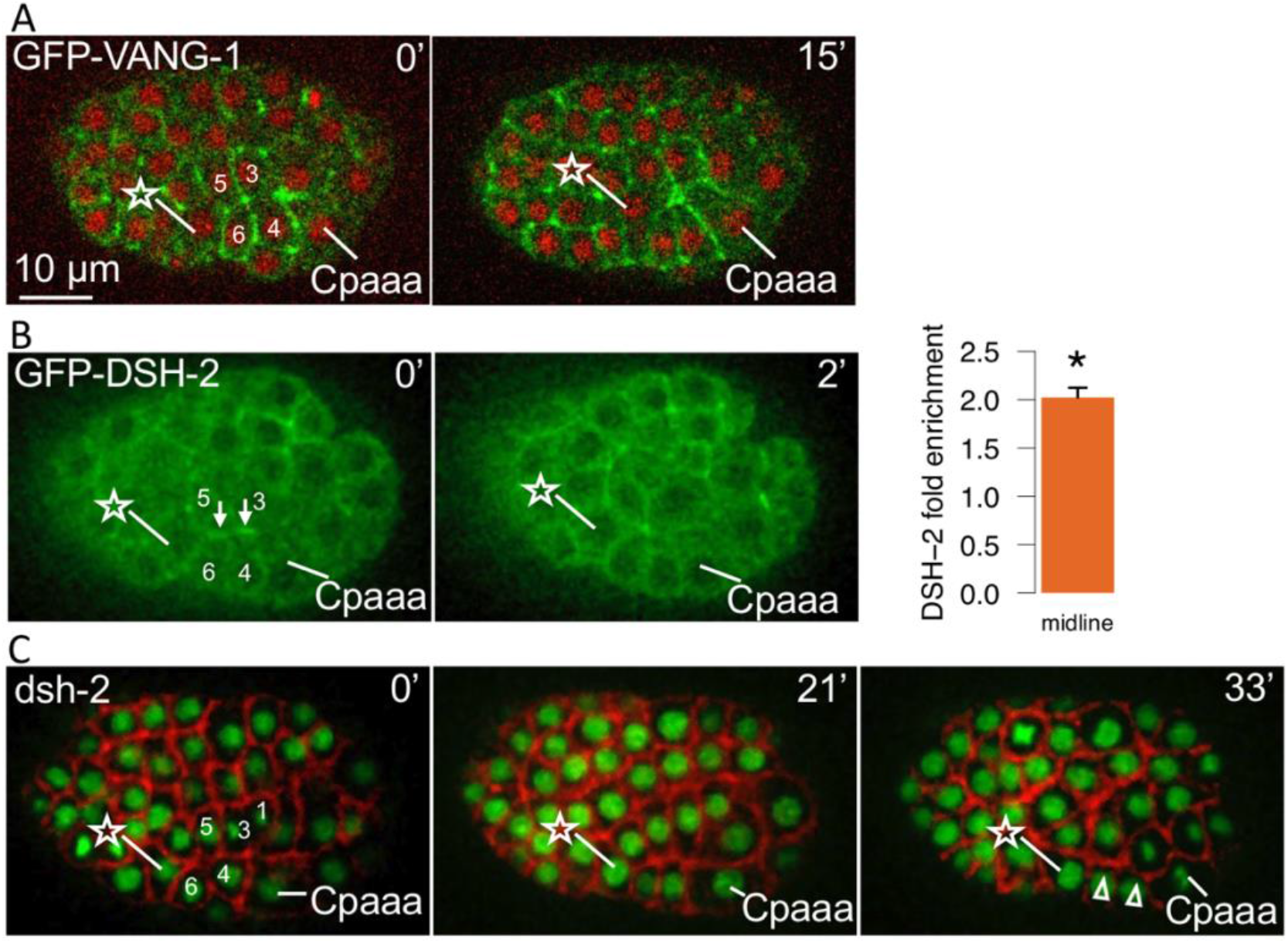
Supporting data for localization of VANG-1 and MIG-1. (A) GFP-VANG-1 in Cpaaa sequential rosettes in another wildtype embryo. See Figure 4A for convention. (B) CRISPR/GFP-DSH-2 (*cp51*) in sequential rosettes of Cpaaa. The arrows denote the midline before first edge contraction. Note nuclei are not labeled in this strain. Bar plots show the fold enrichment of DSH-2 between midline and lateral edges. Error bar, standard error (N=3 embryos). *, Student’s t test p < 0.05. (C) Contraction defects in embryos treated with *dsh-2(RNAi)*. The star and numbers denote 7 ABarp cells involved in the sequential rosettes (see Figure 2). The arrowheads at 33 min denote edges with contraction defect. Nuclei are marked by green channel and plasma membrane is marked by red channel as defined in Figure 1D. All embryos are dorsal view and anterior to the left.

**Table S1.** Genetic interactions of *vang-1(ok1142), mig-1(e1787)*, and *lat-1(RNAi)* knockdown. Fisher’s exact test on contraction defects (top) and resolution defects (bottom) between double loss vs. single loss and triple loss vs. double loss. See Table 1 for counts of phenotypes in each mutant. In double loss vs. single loss, only the single loss with stronger phenotype was compared.

| Contraction defects |  | P value of Fisher's exact test |
| --- | --- | --- |
| *mig-1, lat-1(RNAi)* **vs.** *lat-1(RNAi)* |  | 0.61 |
| *vang-1, lat-1(RNAi)* **vs.** *lat-1(RNAi)* |  | 0.45 |
| *mig-1, vang-1* **vs.** *mig-1* |  | 1 |
| *mig-1, vang-1, lat-1(RNAi)* **vs.** | *mig-1, lat-1(RNAi)* | 0.0087 |
|  | *vang-1, lat-1(RNAi)* | 0.0041 |
| | *mig-1, vang-1* | $2.9 \times 10^{-7}$ |

**Table S1. Genetic interactions of *vang-1(ok1142)*, *mig-1(e1787)*, and *lat-1(RNAi)* knockdown.**
| Resolution defects |  | P value of Fisher's exact test |
| --- | --- | --- |
| *mig-1, lat-1(RNAi)* **vs.** *mig-1* |  | 1 |
| *vang-1, lat-1(RNAi)* **vs.** *vang-1* |  | 1 |
| *mig-1, vang-1* **vs.** *vang-1* |  | 0.58 |
| *mig-1, vang-1, lat-1(RNAi)* **vs.** | *mig-1, lat-1(RNAi)* | 0.46 |
|  | *vang-1, lat-1(RNAi)* | 1 |
|  | *mig-1, vang-1* | 1 |

## Acknowledgements

We thank Dr. Jennifer Zallen for discussions. This work is supported by NIH R01GM097576 to Z.B. and an NIH center grant to MSKCC (P30CA008748).

## Author Contributions

Y.X. and Z.B. conceived the project. Y.X., Y.C. and A.C. performed experiments and data analysis. Y.X. and Z.B. wrote the manuscript.

## MATERIALS and METHODS

### 1. *C. elegans* strains and genetics

*C. elegans* were cultured at room temperature on NGM plates with OP50 bacteria as previously described (Brenner, 1974). N2 Bristol was used as the wildtype strain. See Table S2 for all strains used in this study. Genetic crosses were performed according to standard protocol (Fay, 2006). RNAi experiments were performed according to standard feeding protocol using RNAi clones from commercially available RNAi libraries (Du et al., 2015).

**Table S2.** *C. elegans* strains used in this study.

| Designation | Source | Identifiers | Note |
| --- | --- | --- | --- |
| *ItIs44 [pie-1p::mCherry::PH(PLC1delta1) + unc-119(+)] zuls178 [his-72p (1kb)::his-72::SRPVAT::GFP::his-72 3UTR + unc-119(+)] V* | (Pohl and Bao, 2010) | *BV24* | Nuclei and cell membrane |
| *mig-1(e1787) I* | Caenorhabditis Genetics Center | *CB3303* | *mig-1* mutant |
| *vang-1(ok1142) X* | Caenorhabditis Genetics Center | *RB1125* | *vang-1* mutant |
| *Ex[Pmig-1::MIG-1::GFP]* | This study | *BV600* | MIG-1 localization |
| *zy60[GFP::VANG-1] X* | (Shah et al., 2017a) | *OU410* | VANG-1 localization |
| *nmy-2(cp13) I* | Caenorhabditis Genetics Center | *LP162* | NMY-2 localization |
| *cp51[mNeonGreen::dsh-2] II* | (Heppert et al., 2018) | *LP228* | DSH-2 localization |
| *sax-3(ky123) X* | Caenorhabditis Genetics Center | *CX3198* | *sax-3* mutant |
| *vab-1(dx31) II* | Caenorhabditis Genetics Center | *CZ337* | *vab-1* mutant |
| *tol-1(nr2013) I* | Caenorhabditis Genetics Center | *IG130* | *tol-1* mutant |

### 2. Plasmid construction of Pmig-1:MIG-1:GFP fusion

A 3.9 kb *mig-1* promoter (Mizumoto and Shen, 2013) and *mig-1* isoform a cDNA was cloned into BamHI/KpnI sites pPD95.75 by Gibson Assembly. All amplified fragments were verified by sequencing. Primers sequences: 5’-AAGCTTGCATGCCTGCAGGTCGACTCTAGAGGATCC, 3’-GGTACCGGTAGAAAAAATGAGTA.

### 3. Embryonic imaging

Live imaging was performed as previously described (Bao and Murray, 2011). In brief, ∼5 gravid adult worms were picked to M9 buffer drop (3 g KH2PO4, 6 g Na2HPO4, 5 g NaCl, 1 ml 1 M MgSO4, per liter H2O) to wash away OP50 bacteria. The worms were then transferred to a second drop of ∼20 µl M9 buffer and cut in the middle to release embryos. 10-20 embryos at 2-4 cell stage were transferred to 1.5 µl M9 buffer mixed with 20 µm polystyrene beads on a 24 × 50 mm coverslip in and sealed with vaseline under an 18 × 18 mm smaller coverslip. The following setup of 3D time-lapse imaging was dependent on the purpose of experiment.

For examining the movement phenotype of Cpaaa with nuclei and plasma membrane labeling, ∼20 embryos were transferred on the coverslip and arranged into 2-4 embryos per microscopy stage. Images were acquired on a spinning-disk confocal microscope and Zeiss Observer Z1 microscopy with Zeiss PlanApo 40x/1.3 Oil objective. Embryos were imaged using 30 focal planes spaced 1 µm apart at 20 °C at 60 seconds intervals for 4 hours.

For examining the protein localization with GFP during Cpaaa movement, ∼10 embryos were transferred on the coverslip and arranged into 1-2 embryos per microscopy stage. Images were acquired on a spinning-disk confocal microscope comprising a Zeiss Axio Observer Z1 frame with an Olympus UPLSAPO 60XS objective, a Yokogawa CSU-X1 spinning-disk unit, and two Hamamatsu C9100-13 EM-CCD cameras. Embryos were imaged using 30 focal planes spaced 1 µm apart at 20 °C at 75 seconds intervals for 4 hours.

### 4. Image analysis

a. Lineage analysis Automated cell lineage tracing was implemented and curated in StarryNite and AceTree pipeline (Bao et al., 2006; Boyle et al., 2006). For embryos used for screening of cell migration, the complete lineage was curated until the 350-cell stage. For embryos used for examining movement phenotype of Cpaaa and protein localization around Cpaaa rosettes, the lineages of Cpaaa and 7 ABarp cells were curated until the cell division of Cpaaa.
b. Intensity measurement The intensity measurement of GFP signal was performed in Fiji software as previously described (Shah et al., 2017a). For a given cell-cell edge, a 3 pixel-wide line segment tool in Fiji was manually applied on the edge to measure the intensity. The mean per-pixel intensity was defined as the signal intensity on the edge after background subtraction, which was the median of a 75×75 pixel square in cell cytoplasm. The mean of signal intensity for each set of edges was calculated in each embryo. Student’s t-test was applied to test the fold difference between edges of interest and control edges in N≥3 embryos, i.e., vertical vs. horizontal, midline vs. lateral, vertical vs. lateral.

### 5. Computational screen of large neighborhood change

a. Embryonic stages based on division timings Most cell divisions are synchronous within each founder lineage of *C. elegans* embryos and are progressively prolonged (Bao et al., 2008). The cell size and lifetime have a dramatic difference from 1-cell to 350-cell (end of gastrulation), thus it is necessary to divide this period into stages. To this end, we searched for timepoints when massive parallel cell divisions are just finished as the boundary between stages (Fig. S2A). First, local maximum on the increase of cell number between t_i_ and t_i-1_ (D_i_) was defined by D_i_ > 5 and D_i_ > D_j_, for j ∀ [i-10, i+10]. Second, the boundary between stages was defined as the first timepoint after D_i_ with D = 0. Four boundaries were found that give 5 stages. The first boundary corresponds to the start of gastrulation (26-cell stage), indicating the definition of boundaries is rational. We found cells in stages III and IV have significantly larger migration distance than cells in phase I (Fig. S2B-C, adjusted p value = 3×10^−7^ and 4×10^−5^ by Mann-Whitney U test, respectively). Thus, we chose cells in stages III and IV to search for candidate cells of large neighborhood change.
b. Large neighborhood change To search for large neighborhood change in candidate cells, we considered neighborhood change to exclude whole-embryo rotation. The neighborhood change for each cell is defined as the average of distances to initial neighbors when the cell is dividing and distances to final neighbors when the cell is born (Fig. 1A). Cells with neighborhood change above threshold mean + standard deviation were selected and examined in images of embryos with fluorescently labeled nuclei and plasma membrane for potential T1 process, rosette, and sequential rosettes.

